# Does learning to read change the perception of speech? Evidence from perceptual recalibration of speech sounds in adult illiterates, semi-literates, and literates of Tamil

**DOI:** 10.1101/2021.08.27.457915

**Authors:** Holger Mitterer, Mrudula Arunkumar, Jeroen van Paridon, Falk Huettig

## Abstract

How do different levels of representation interact in the mind? Key evidence for answering this question comes from experimental work that investigates the influence of knowledge of written language on spoken language processing. Here we tested whether learning orthographic representations (through reading) influences pre-lexical phonological representations in spoken-word recognition using a perceptual learning paradigm. Perceptual learning is well suited to reveal differences in pre-lexical representations that might be caused by learning to read because it requires the functional use of pre-lexical representations in order to generalize a learning experience. In a large-scale behavioural study in Chennai, India, 97 native speakers of Tamil with varying reading experience (from completely illiterate to highly literate) participated. In marked contrast to their performance in other cognitive tasks, even completely illiterate participants showed a perceptual learning effect that was not moderated by reading experience. This finding suggests that pre-lexical phonological representations are not substantially changed by learning to read and thus poses important constraints for the debate about the degree of interactivity between different levels of representations during human information processing.

## Introduction

The nature of the interaction between different levels of representations during human information processing is of crucial importance for our understanding of the human mind and brain. The notion of strong modularity between processing levels (Fodor, 1983) has fallen out of favour with large parts of the cognitive science community in recent years. The *extent* of interactivity between different types of representations and information-processing streams, however, remains hotly debated. An important testing ground to assess the interactivity of the human mind is the nature of the interplay between orthographic and phonological representations. The present study addresses this question by investigating whether orthographic knowledge restructures pre-lexical phonological representations as people learn to read. We define pre-lexical phonological representations here as intermediate units that are activated by auditory input and, in turn, lead to activation of word candidates, such as the phoneme units in the TRACE model of spoken-word recognition (McClelland & Elman, 1986).

Written language is one of the most significant cultural inventions in human history (Huettig, Kolinsky, & Lachmann, 2018; Morais, 2018). Reading and writing has had a huge impact on society; modern societies are almost unimaginable without the ability to “store” knowledge in written language. Written knowledge, however, does not only influence human behaviour on a societal but also on an individual level, as people learn to read and write. Learning to read and write improves mirror image discrimination (Fernandes et al., 2021; Kolinsky et al., 2011; Pegado et al., 2014), face recognition (Van Paridon et al., 2021), visual search (Olivers et al., 2014), verbal memory (Demoulin & Kolinsky, 2016; Smalle et al., 2019), prediction during spoken language processing (Favier, Meyer, & Huettig in press; Huettig & Pickering, 2019; Mishra et al., 2012), and even the perception of facial emotions (Eviatar, 1997) and non-verbal intelligence as measured by Raven’s progressive matrices (Hervais-Adelman et al., 2019; Olivers et al., 2014; Skeide et al., 2017). Given this large impact of literacy on cognitive processing it is reasonable to assume that the acquisition of orthographic representations may also influence speech perception.

### Phonological awareness & thinking about speech

Indeed, it has been conclusively demonstrated that acquiring an alphabetic script changes phonological awareness, i.e. our *thinking* about speech (Lukatela, et al., 1995; Morais, Cary, Alegria, & Bertelson, 1979; Morais et al., 1986; Prakash et al., 1993). Some phonological awareness can occur without instruction (e.g., noticing words that rhyme, or sound repetition as in “Susie sold six salami sandwiches”) but pre-literate children typically do not succeed when asked to delete the sound “b” from “bit” (resulting in “it”) or substituting “c” in “cat” with “h” (resulting in “hat”). Morais and colleagues (Morais et al., 1979; Morais, Content, Cary, Mehler, & Segui, 1989) showed in a series of experiments that this is not a simple developmental difference between children and adults. Illiterate adults who, for socio-economic reasons, have not learned to read are also unable to perform successfully in tasks that require the addition, deletion, or addition of a phoneme to a given word. Phonemic awareness usually does not arise without formal reading instruction in an alphabetic script (Kolinsky & Morais, 2018). In other words, learning to read changes how we *think* about spoken language. This raises the question of whether learning to read also changes how we *perceive* spoken language.

### Segment-sized units and the perception of speech

Most models of spoken-word recognition assume that we recognize words as being made up of smaller segments such as phonemes (e.g., McClelland & Elman, 1986; Norris, 1994). The finding that we only become aware of phonemes when learning to read however raises the question of whether such small segment-sized units are actually ‘used’ in spoken-word recognition. Indeed, some researchers (e.g., Port, 2007), argue that segment-sized units are not used at all in spoken-word recognition. Port (2007), for example, suggested that alphabetic scripts had undue influence on the field of phonology, and—by extension—on cognitive science itself, by “tricking” scientists into believing that speech processing makes use of letter-sized units. In a similar vein, Goldinger (1998) proposed a model of spoken-word recognition that assumed that words were recognized by comparing them holistically to episodes of previous encounters of the same word, a strategy called “lexical access from spectra”, somewhat similar to the model proposed by Klatt (1979). However, these proposals have remained controversial. Klatt himself (1989) noted that it is difficult to come up with a similarity metric that is able to discard irrelevant spectral differences—like those between speakers with different vocal-tract sizes—but make use of all the available fine-grained information. There is a broad consensus that there is some form of intermediate unit in the spoken modality, akin to the letters in written language (Cutler et al., 2010; Fowler et al., 2016; McQueen et al., 2006; Mitterer et al., 2013).

### Reading-induced phonological restructuring

If there are some segment-sized units that are functional in spoken-word recognition, then learning to read might influence spoken-word recognition by changing the phonological representations of words. According to such an account, orthographic experience directly shapes the representational structure of the spoken word-recognition system. There is some evidence that this can occur on the lexical level. Experience with a new written word, for instance, influences lexical processing in the spoken modality, most likely through the generation of a lexical entry in the auditory modality (Bakker et al., 2014). Phonological restructuring however implies an influence of orthographic experience on the pre-lexical level, so that pre-lexical representations in the auditory modality are influenced by learning to read. One piece of evidence consistent with such a view comes from an eye-tracking study comparing Hindi-speaking literates and illiterates in India. Huettig, Singh, and Mishra (2011) administered a “look and listen” eye-tracking task in which participants were viewing a screen with four objects while listening to a simple sentence. Participants were asked to listen to the sentences and not to take their eyes off the screen. On the critical trials, the critical word in the sentence (e.g., *crocodile* in “They saw a crocodile”) was not shown on the screen. Instead, the screen depicted a semantic competitor (e.g., *turtle*) and a phonological competitor (e.g., *peas*; in Hindi, the phonological forms of *crocodile* and *peas* share their three initial phonemes). Literate participants showed a clear peak in looks to the phonological competitor when hearing the (identical) first syllable, and then looked more and more to the semantic competitor (as in similar previous experiments, e.g., Huettig & McQueen, 2007). Illiterates, however, never looked more at the phonological competitor than at unrelated distractors. Instead, they paid more attention to the semantic competitor. In a second experiment, Huettig et al. (2011) tried to boost looks to the phonological competitor by replacing the semantic competitor with an unrelated distractor. The literate participants’ pattern of looks to the phonological competitor was hardly influenced by this change; there still was a clearly defined peak of looks to the phonological competitor that was time-locked to the phonological overlap between the competitor name and the spoken word. Illiterates, however, were now also influenced by phonological relatedness, but their looks did not show a clear peak that was time-locked to the phonological overlap. Instead they looked somewhat more at the phonological competitor during most of the trial rather than showing a clear, but time-locked (and time-limited) peak as the readers did. This finding is consistent with evidence from neuroimaging studies that found different patterns of activation for literate and illiterate participants in the planum temporale, a site linked to auditory language processing, during speech comprehension (cf., Dehaene et al., 2010, 2015; but it is noteworthy that Hervais-Adelman et al., 2021, failed to replicate such an effect with a similar population, albeit with a different language and writing system). One way to explain these results is by appealing to the notion that the phonological representations of illiterates are less well defined or less segmental and more holistic.

This reasoning dovetails well with the claim that early word-form representations are holistic and only become more segmental based on experience with a larger number of words. An underlying idea here is that, for a small vocabulary, holistic recognition is sufficient; only when vocabulary size grows is it necessary to generate segmental representations to distinguish more and more similar sounding words. This has become known as the lexical restructuring hypothesis (Metsala & Walley, 1998). Because vocabulary size is boosted by print exposure (Hayes, 1988; Stanovich & Cunningham, 1992), it is conceivable that literacy may strongly boost lexical restructuring. This assumption would explain the findings by Huettig et al. (2011) that illiterates seem to match phonological information more globally and not in such a fine-grained manner as literates.

There is however also evidence that appears to be (at least) inconsistent with the notion that learning to read results in the restructuring of lexical representations. Investigations employing low-level perception tasks (e.g., categorical perception or phonetic fusion in dichotic listening), for instance, mostly did not find clear differences between literate and illiterate adults (for a review, see Morais & Kolinsky, 2019). Kolinsky and colleagues for example argue that reading acquisition does not change categorical perception of speech sounds, though they propose that literacy exerts its influence on categorical precision. Kolinsky et al. (2021) performed identification and discriminations task with a voicing continuum (/de/-/te/) with Brazilian literate and illiterate adults as well as Grade 2 children (around 8 years of age). The results showed that discrimination performance was well predicted by identification performance in all groups. It is important to point out here that the notion of categorical perception as a hallmark of speech perception is currently questioned by a number of influential researchers in the field of spoken-word recognition (Brown-Schmidt & Toscano, 2017; McMurray et al., 2009; Schouten et al., 2003; Toscano et al., 2010). Given the controversy about the relevance of categorical perception for speech perception, the more interesting and crucial result of the Kolinsky et al. (2021) study is that illiterates and (still) low-literate children showed a less steep category boundary than literate adults. A literacy-induced steeper category boundary suggests that reading experience leads to more precise *pre-lexical* phonological representations. The Kolinsky et al. (2021) study therefore can be interpreted as providing support for the notion of reading-induced restructuring of phonological representations. In the present study we have a fresh look at this conclusion by employing the perceptual-learning paradigm with adult illiterates, semi-literates, and literates of Tamil in Chennai, India.

### Perceptual learning as a window on pre-lexical representations

The perceptual learning paradigm (Norris et al., 2003) has been found to be particularly informative about the structure of pre-lexical phonological representations. In this paradigm, participants go through an exposure phase, followed by a test phase. During exposure, participants are exposed to ambiguous speech sounds in unambiguous contexts. In most early experiments on the effect, the ambiguous speech sound was an ambiguous sound between /f/ and /s/ (henceforth transcribed as [^s^/_f_]). This ambiguous sound then is used to replace /s/ and /f/ in words ending on /s/ or /f/ (e.g., *giraffe*: /ʤəɹaf/ → [ʤəɹa^s^/_f_] or *platypus*: /plætəpʊs/ → [plætəpʊ ^s^/_f_]. How this replacement occurs is varied between participants: Half of the participants hear the /f/-words with the edited fricative [^s^/_f_] and /s/-words with a normal [s], while the other half hear the /s/-words with the edited fricative and unedited /f/-words. Because minimal pairs (such as *knife* – *nice*) are avoided, participants can infer the identity of the fricative using lexical knowledge, analogue to the Ganong (1980) effect. Indeed, in versions using a lexical-decision task during exposure, most participants identify the edited words as valid words most of the time (see, e.g., Mitterer & Reinisch, 2013; Norris et al., 2003). The important question here was whether lexical knowledge is not only used to identify the intended words but also re-calibrate the perception of the /s/-/f/ contrast. This is then tested in the test phase, during which participants are asked to identify tokens along an /s/-/f/ continuum (e.g., /εs/-/εf/). In line with the assumption of re-calibration, participants who heard the ambiguous fricative in /s/-final words more often identify tokens from the middle of the continuum as /s/ in comparison to participants that heard the ambiguous fricative in /f/-final words (for a review, see Samuel & Kraljic, 2009).

The perceptual-learning paradigm is particularly interesting for the current research question because it shows *functional generalization* used to overcome the invariance problem in speech perception (Mitterer & McQueen, 2009). Much work supports the notion that the recalibration affects pre-lexical phonological representations. There are two types of evidence supporting this conclusion. First, learning generalizes from the exposed words to other words (Mitterer et al., 2011), and this generalization also occurs when the measure is implicit (e.g. cross-modal priming, see McQueen et al., 2006; Sjerps & McQueen, 2010). Second, evidence from eye-tracking (Mitterer & Reinisch, 2013) suggests that recalibration affects perception early on during speech perception, roughly at the same time that acoustic differences are processed. Taken together this strongly suggest that perceptual learning reflects pre-lexical phonological representations that intervene between acoustic input and the mental lexicon. We therefore consider perceptual learning to be an exceptionally strong paradigm for revealing potential influences of learning to read on the structure of phonological representations.

### Perceptual learning and reading acquisition

Let us consider how reading experience could influence perceptual learning. Such a prediction follows directly from the claim—as made by Kolinsky et al. (2021)—that reading experience leads to better defined pre-lexical phonological representations. Note that perceptual learning relies on the perceiver to “notice”—the quotation marks here are meant to indicate that it is not a conscious noticing—a difference between the modified, ambiguous input and the pre-lexical phonological representation. However, (literacy-related) less precise specification of those pre-lexical representations will mean that the modified input is still considered within the range of acceptable inputs for the pre-lexical category and hence should fail to trigger learning. Interestingly, such a prediction from the Kolinsky et al. (2021) results is in contrast with the findings of a study by McQueen, Tyler, and Cutler (2012), who used the perceptual learning paradigm with a group of pre-literate and literate children (aged around 6 and 12, respectively). In order to make the procedure more child-friendly, they made two changes. First, instead of a lexical decision task for the exposure phase, a picture verification task was used. This allows a wider range of words to be used, including minimal pair words (for instance, *knife*, disambiguated by an accompanying picture, so that listeners can rule out that the intended word would be *nice*) and words with the critical fricative in word-initial position (e.g., *flamingo*)^1^. Secondly, the test phase made use of a newly learned minimal pair: two puppets from a 1980’s children program (*Fraggle Rock*) were introduced as *Simpie* and *Fimpie*. Participants then heard these words with tokens from a fricative continuum and had to identify the “correct” puppet. Six-year olds turned out to be able to do this task and showed a similar shift in their identification responses based on the exposure as adults and 12-year olds.

The McQueen et al. (2012) findings thus strongly suggests that 6-year olds already have adequate pre-lexical phonological representations on par with older listeners and do not provide any support for literacy-induced phonological restructuring. There are however three caveats that caution strong conclusions. First, it is conceivable that the same behavioural pattern by 6- and 12-year olds may come about by different underlying mechanisms. Less well specified pre-lexical representations, leading to smaller learning effects, may be counteracted by greater flexibility of these categories. While this may seem far-fetched, it becomes plausible when one considers that the average ability for phonological acquisition of a second language declines strongly between 6 and 12 years (e.g., Flege et al., 2006). Second, the choice of the fricatives /s/ and /f/ may also overestimate the fidelity of pre-lexical representations, since /s/ and /f/ are segments that are easy to acquire, as they stand out of the speech stream rather clearly. For instance, Remez, Rubin, Berns, Pardo, and Lang (1994) have argued that it is in fact a challenge to keep the speech stream “together” rather than “streaming out” with voiceless fricatives, such as /s/ and /f/, which contain mostly high-frequent aperiodic acoustic energy from the rest of the speech signal that is mostly found below 3kHz and is mostly periodic. Moreover, voiceless fricatives are relatively stable over syllable positions, and learning can occur across syllable positions for fricatives (Jesse & McQueen, 2011) but not for segments that vary more strongly acoustically over syllable positions (Mitterer et al., 2013; Mitterer & Reinisch, 2017). Finally, the six-year old participants in Australia in McQueen et al. (2012) most likely had some experience with letter names, for instance through the letter naming games that are part of the Australian pre-school curriculum. As such, it remains a possible despite the data provided by McQueen et al. (2012) that learning to read (or letter knowledge) may influence the fidelity of pre-lexical phonological representations.

### The present study

The inconsistent general findings of the influence of learning to read on the structure of phonological representations and the caveats concerning the interpretation of McQueen et al. (2012) thus warrant another careful examination of this issue. To this end, we tested perceptual learning in adult literates and illiterates rather than young and older children (thereby avoiding age and developmental confounds in the group comparison of readers and pre-readers). Literate and illiterate Tamil speakers in Chennai, India, who were carefully matched for socioeconomic variables took part in the study. Chennai is the sixth largest city in India, a metropolis with an illiterate population of about 10% (Census, 2011). The common factors for illiteracy are socioeconomic: poverty and family situations result in a substantial number of neurologically normally developed people who did not attend any formal schooling and hence do not know to read or write Tamil.

In order to provide a strong test of pre-lexical representations we used the dental versus retroflex stop contrast as the target for recalibration and tested three socio-economically similar groups of individuals with differing reading ability. This is a contrast that typically is difficult to perceive for non-native speakers (Werker & Tees, 1984), and as a contrast between stop consonants, is more reliant on dynamic information and less obviously “segmental” than the contrast between voiceless fricatives used by McQueen et al. (2012).

Reading experience is more likely to lead to a more segmental perception if the orthography supplies cues to those segments. Tamil uses an “akshara-based” orthography and because such orthographies have sometimes wrongly been described as syllabic (Bhide, 2018), one might question this. All Indic scripts are neither fully alphabetic nor are they syllabic but are best described as an abugida (Daniels, 2021). The features of Indian writing systems are complex and beyond the remit of the current paper but we note here that the Tamil abugida has been described as a “more alphabetic and less syllabic akshara-based orthography” (Bhuvaneshwar & Padakannaya, 2013). Tamil combines basic and separable shapes for each segment into akshara blocks. Each consonant has a basic shape which remains invariable in combination with different vowels. These properties can be exemplified well with the minimal pair used in this study: /kad̪aɪ/versus /kaɖaɪ/ (English: ‘story’ vs. ‘shop’). These words are written as 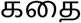 versus 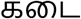 The first grapheme 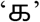 stands for the syllable /ka/, derived from the grapheme 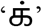 for /k/, and the removal of the diacritic dot indicates the combination with the consonant /a/. The diphthong /aι/, which is pronounced as [εi] in non-initial position, is represented by the symbol 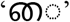 which always precedes its onset consonant. Those are represented by 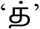 for the dental /d̪/ and 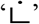 for the retroflex stop /ɖ/ as single letters and combine to 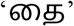 and 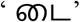 for /d̪aɪ/ and /ɖaɪ/. The important point is that although Tamil script is quite different from an alphabetic script such as English, it contains nevertheless distinct and clear cues to segmental perception.

Three groups of participants took part in our study. The first group were illiterates with no formal schooling and who had no knowledge about reading and writing. Participants who could read and write were divided into two groups: one comprising those who completed secondary education (literates) and the other group consisting of those who attended formal schooling but dropped out or discontinued during school (semi-literates). On average, the literates had completed 11 years of schooling whereas semi-literates had completed 6 years of schooling.

Based on the above considerations, the predictions are straightforward: If segmental representations evolve independent of reading experience, all groups should show similar amounts of recalibration. If pre-readers, however, do not make use of segmental representations, but more holistic units, recalibration should only be observed with the groups that have reading experience.

## Method

### Participants

97 native speakers of Tamil participated in the experiment. Three participants were excluded from further analysis because they were outside the specified age range of 25 to 45. Forty-seven of the remaining participants were female. Participants fell into one of three groups of reading ability. Thirty participants were proficient readers, 30 had some reading ability but were not fully literate, and 34 had never been taught to read (see control tasks in results section for further information about participant characteristics).

### Stimuli and Apparatus

Experiments were performed at ARUWE, (a non-governmental organization (NGO) working towards enhancing the lives of the urban poor living in Chennai living by providing quality care, support, advocacy, information, education, research, training and development. Experiments were run on a Windows-operated laptop using PsychoPy (Peirce et al., 2019). Sounds were played through a Sennheiser HD205 closed headphone. Participants responded by pressing keys marked with a green sticker (left) or red sticker (right) on the experiment laptop.

Perceptual-learning experiments require a larger set of exposure stimuli containing one member of the chosen phonological contrast (i.e., dental d̪/ vs. retroflex /ɖ) in words that do not form minimal pairs based on that contrast and at least one minimal pair of that same contrast for the test phase. For the exposure phase, we made use of a picture-verification task with 104 trials. For this task, we found 22 words each with a dental and a retroflex stop in intervocalic position, for which matching pictures could be found, leading to 44 critical exposure trials. For these words, exchanging the dental or retroflex stop led to a nonword. For example, /mʊd̪əl/ was used for the dental exposure (with */mʊɖəl/ being a nonword) and /ʋiːɖʉ/ was used for the retroflex exposure (with */ʋiːd̪ʉ/ being a nonword). Additionally, we found 60 words that were depictable that did not contain a retroflex or dental stop like /pu:nəj/. Forty-three of these were presented with matching picture and 17 with mismatching pictures. For the test phase, we used the minimal pair /kad̪aɪ/ versus /kaɖaɪ/ (English: ‘story’ vs. ‘shop’).

For all words, pictures were gathered using a Google image search. It was not always possible to find pictures that matched a word to the extent that it would elicit the target word in a picture-naming task (e.g., for the critical word *story* an open book with some animated characters emanating from it was used). Note, however, that participants only need to say that there is a match between picture and spoken word, which does not imply that the image by itself activates the intended word only.

We calculated the overlap between exposure and test items for both diphones involving the critical stop. The exposure items with a dental stop contained two examples each of both the VC and the CV diphones and in the retroflex set, there were six items with the same VC diphone and three with the same CV diphone. This is well below the threshold of the ten items that would be necessary for learning, so that successful learning can only be achieved by segment-sized representations.

To generate sound files for the exposure phase, we recorded a 45-year old female native speaker of Tamil, using the Speechrecorder program (Draxler & Jänsch, 2004). The fillers were recorded once, but the critical items for exposure and test were recorded twice. For the critical exposure items, the words were also recorded with the critical segment exchanged—that is, with a retroflex where there should be a dental stop, and vice versa—to allow us to generate an audio-morphing continuum between the real words and their non-word counterparts containing the other segment.

The recordings were further processed by trimming initial and final silences and finding segment boundaries. These segment-boundary timings were then used in the time-aligned version of the Straight audio-morphing algorithm (Kawahara et al., 1999). This morphing algorithm allows us to generate stimuli that are ambiguous between retroflex and dental by morphing the recordings of the words and corresponding nonwords (e.g., /mʊd̪əl/ and /mʊɖəl/). The morphing algorithm provides a source-filter separation of the sound and then interpolates the values for the source and the filter. The interpolated source and filter are then used to generate a new, morphed sound.

Audio continua were generated with 11 steps from a fully retroflex re-synthesis to a fully dental re-synthesis, in steps of 10%. Two native speakers of Tamil then decided for each continuum which sounds were perceptually most ambiguous (ranging from 40% to 70% retroflex signal in the mix). These sounds were then used as ambiguous sounds during the exposure phase. From the 11-step test continuum, we used steps 2, 3, 4, 5, 6, 8 and 10 to ensure that all participants would have near-endpoint stimuli to guide them to use both answer options during the test phase, but they would also have exposure to ambiguous stimuli which are likely to be affected by perceptual learning (see Mitterer et al., 2018, for the importance of having variation in the test phase).

### Procedure

Experiments started with the instruction for the exposure phase. Instructions were supplied verbally by the 2^nd^ author, who is a native speaker of Tamil. While this may give rise to slight variations in the instructions, providing controlled written instructions was not possible given that some participants were not able to read.

The experimenter told participants that they would see a picture and hear a word, and should then indicate via a button press whether they matched or not. On each trial of the exposure phase, participants first saw a fixation cross for 400 ms, then a blank screen for 200 ms and a picture plus a “reminder” that indicated that the green (left) button should be pressed if there is a match and the red (right) button if there is a mismatch. This was achieved by presenting, on the left and right of the screen, a thumbs up versus thumbs down icon with additional color coding (green/red for match/mismatch). Two hundred milliseconds after the presentation of the picture, the sound file started playing.

There were 104 exposure trials, of which 60 were fillers. These contained 17 mismatch trials so that participants would be using both answer options during exposure. Different random orders were generated for all participants, with two constraints. For each participant, the experiment started with three filler trials with a match. The rest of the trials were then randomized with the constraint that each trial with an ambiguous speech sound was followed by a clear match. This is in line with earlier studies on perceptual learning of speech sound (Norris et al., 2003) and done to avoid making the manipulations too obvious. Within each group of participants, half of the participants received exposure with the ambiguous sound used instead of a retroflex (ambiguous to retroflex, abbreviated as “amb2rtrflx”) while the other half received exposure with the ambiguous sound used instead of a dental (ambiguous to dental, abbreviated as “amb2dental”).

After the exposure task, participants were instructed that they now would hear a speaker saying either /kad̪aɪ/ (English: ‘story’) or /kaɖaɪ/ (English: ‘shop’) and their task was to decide what the speaker actually said (i.e., two-alternative forced choice (2AFC) task). Two pictures reminded participants which button (green/left vs. red/right) went with what option. The seven stimuli of the test continuum were repeated eight times, with a new randomization on each rollover.

In addition to this experiment, control tasks were also administered where the participants’ reading ability, working memory and intelligence were measured by means of a reading task, a digit span task, and a shortened Raven’s progressive matrices task, respectively.

## Results

### Control Tasks

Participants’ word and pseudoword reading abilities were assessed. As is standard practice in research on reading development (see, e.g., Vandermosten et al., 2020), we administered test that measure how many words (see, .e.g. Brus & Voeten, 1973) or pseudoword (Bos et al., 1994) can be read within a given time (usually one or two minutes). Since such tests are not available for Tamil, a native speaker of Tamil (the 2^nd^ author) chose Tamil words of varying complexity. High Frequent Tamil words ranging from monosyllabic to words containing six syllables were selected and arranged in increasing order of syllable length. The pseudowords were created by manipulating vowel/consonant from a Tamil word such that the resulting pseudowords were phonotactically legal. These also ranged from monosyllabic to pseudowords containing six syllables. Participants were given 60 seconds to read up to 100 items from a list for both words and pseudowords. Response were scored by a native speaker (M. Arunkumar). The reading scores (number of words or pseudowords read correctly, out of 100 words or pseudowords, with a one-minute time limit) were calculated across the three groups. Robust differences were found between the literates, semi-literates and illiterates both in the word reading (*F* = 126.1, *p* < .001) as well as in the pseudoword reading tasks (*F* = 98.56,*p* < .001), thus confirming the correct assignment to the specific participant groups.

Participants’ working memory was assessed using forward and backward digit span tasks. The number of correct responses (until producing consecutive errors) were analysed. In the forward digit span task, literates performed better than illiterates (*t* = 5.56,*p* <.001) as well as semi-literates (*t* = 2.88, *p* < 0.05). Similar results were obtained in the backward digit span task where the literates performed better than illiterates (*t* = 5.61, *p* < .001) and semi-literates (*t* = 2.57, *p* < 0.05). The results replicate studies with similar populations, which found that acquiring literacy results in increased verbal working memory (Demoulin & Kolinsky, 2016; Smalle et al., 2019).

General non-verbal cognitive abilities were assessed using Raven’s progressive matrices. The three groups significantly differ from each other in their performance (*F* = 37.94, *p* < .001). The results replicate those from earlier studies with similar populations, finding that high levels of literacy and education more generally result in robust increases of Raven’s scores (Hervais-Adelman et al., 2019; Olivers et al., 2014; Skeide et al., 2017; though differences in Raven’s between semi-literates and illiterates are typically lower or absent, most likely due to the limited education received by semi-literates). Table 1 shows the list of control tasks done along with the mean and standard deviations of each task across the three groups.

**Table1.** Mean and standard deviation of the control tasks among illiterates, literates and semi-literates.

| Task | Literates |  | Semi-literates |  | Illiterates |  |
| --- | --- | --- | --- | --- | --- | --- |
|  | Mean | SD | Mean | SD | Mean | SD |
| **Word Reading (score out of 100)** | 59.8 | 17.4 | 38.1 | 18.8 | 3.41 | 6.03 |
| **Pseudoword Reading (score out of 100)** | 38.6 | 14.3 | 21.1 | 13.7 | 0.73 | 1.73 |
| **Forward Digit Span** | 7.3 | 1.21 | 5.63 | 1.69 | 4.41 | 2.09 |
| **Backward Digit Span** | 5.17 | 1.66 | 3.67 | 2.40 | 2.21 | 1.89 |
| **Raven's Progressive Matrices (Accuracy in %)** | 32.3 | 46.8 | 19.4 | 39.6 | 12.1 | 32.6 |

We used these results to inform our contrast coding for the group variable. The results from the reading tests show that the literate group read nearly two times as many words and non-words as the semi-illiterate group, but that the semi-illiterates read about *ten* times as many words and non-words than the illiterate group. Therefore, the three-level variable was used to generate one contrast between the two literature groups with the literate group and a second contrast between the two literate groups.

### Exposure Phase

Figure 1 shows how often the participants accepted the pictures as matching the presented word. Though these data are not critical for our theoretical question, it is necessary to show that, first, the participants understood the task and, second, that there are no large differences in how many of the words with ambiguous stops were accepted as existing words. Sjerps and Reinsich (2015) found that participants who reject more than half of the edited words show reduced or no perceptual recalibration.

**Figure 1.**
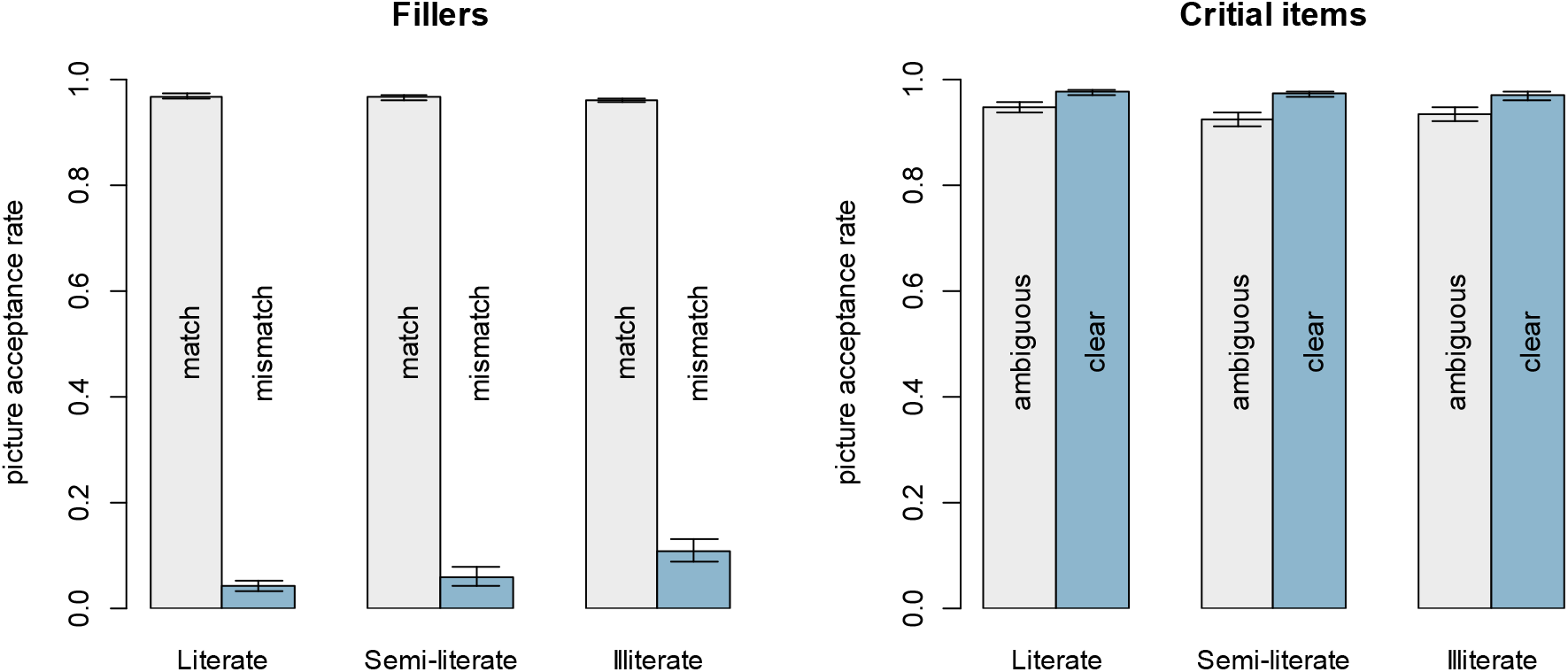
Acceptance rate in the picture-verification task for the three groups for critical items (left panel) and fillers (right panel). Means are based on log-odds of acceptance rates, transformed back into proportions. Errors bars show the standard error based on participants, calculated in log-odds and transformed back into proportions.

The data from the filler trials show that all participants understood the task, accepting matching fillers and mostly rejecting mismatching fillers. For the critical items, we saw slightly lower acceptance rates when the critical word was presented in the ambiguous form (with the stop being ambiguous between dental and retroflex), a pattern also observed in previous studies (Mitterer & Reinisch, 2013). However, even the ambiguous items were accepted in more than 90% of the cases without an obvious difference between the groups (see Figure 1).

Given that it is theoretically important to have an estimate to what extent the data support null-hypotheses, we used Bayesian generalized mixed effect models, using the package *brms* (Bürkner, 2020) in R (version 3.5.3). We used a logit link function (by specifying the Bernoulli family in the brm function) to account for the binary nature of the dependent variable. We specified weakly informative priors with normal distribution with standard deviation of two (i.e., Normal(0,2)) for the contrast-coded predictors that had a range of one^2^. This makes effects of more than four logit units unlikely.

We used model comparisons to estimate the weight of the evidence for the null-hypothesis as a Bayes Factor. To compute this Bayes Factor (BF) via bridge sampling, we drew 10,000 posterior samples per chain after drawing 2000 warm-up samples; running four chains for a total of 40,000 posterior samples (in line with the recommondations by Vasishth et al., 2018). The scripts are available as supplementary materials. We specified model comparisons so that the numerator model was the model with an added effect present and the denominator model had the effect absent. The results BFs are BF10, that is, they specify the evidence for the presence of an effect over the absence of an effect. BFs over 3 are substantial evidence for the presence of an effect and BFs over 10 are strong evidence. For BFs smaller than one, the absence of an effect is more likely than its presence (given the priors), with the inverse values providing the same amount of evidence (i.e., BF < 1/3 means substantial and BF< 1/10 strong evidence).

For the results from the exposure phase, the fixed effects were Ambiguity (clear vs. ambiguous) and Group (i.e., Literate, Semi-Literate, and Illiterate). These fixed effects were contrast coded. For Ambiguity, the ambiguous items were mapped on -0.5 and the clear items on 0.5. For the fixed effect of Group, two orthogonal contrasts were used, one contrasting the two literate groups with the illiterates, and one contrasting full versus semi-literates. The random effects were item and participant. The model made use of a maximal random-effect structure. Table 1 shows the results of the analysis. None of the effects has a credible interval that excludes zero, though the effect of ambiguity has a range that most includes an effect in the direction that ambiguous items are accepted less often. Previous perceptual learning studies found such an effect when participants performed a lexical decision task on the items with an ambiguous fricative (McQueen et al., 2006; Norris et al., 2003) but not with picture matching (McQueen et al., 2012). As such, it may not be surprising to find a small effect here. Most importantly, there is evidence that the groups are likely to behave similarly with regard to the ambiguity, with a BF that is smaller than 0.1. That is, it is unlikely that the groups are differentially affected in how they accepted the ambiguous items compared to the clear items.

**Table 1.** Results from the mixed-effect models on the acceptance rate in the picture-verification task during the exposure phase

|  | B (SE) | 95% CI | Bayes Factor |
| --- | --- | --- | --- |
| Intercept | 4.68 (0.36) | 4.01-5.41 |  |
| Ambiguity | 0.91 (0.51) | -0.07-1.94 | 1.36 |
| Group |  |  |  |
| Literate vs. Illiterate | 0.5 (0.45) | -0.37-1.39 | 0.27 |
| Full Literate vs. Semi-literate | 0.78 (0.54) | -0.26-1.88 |  |
| Group x Ambiguity |  |  |  |
| Literate vs. Illiterate:Ambiguity | 0.20 (0.46) | -0.69-1.10 | 0.08 |
| Full Literate vs. Semi-literate:Ambiguity | -0.19 (0.66) | -1.47-1.13 |  |

Note: Bayes Factors are based on model comparisons with the “fuller” model in the numerator and the reduced model without that effect in the denominator. As such, values above one are indicative of evidence for an effect (with the evidence being substantial above 3 and strong above 10) while values below one indicate more evidence for the null hypothesis.

### Test Phase

Figure 2 shows the descriptive results of the test phase, comparing the two exposure conditions within each group. All groups show a clearly categorical response pattern, with hardly any uncertainty for the two “guiding” stimuli that were mostly dental (0.1 mixRetroflex) or mostly retroflex (0.9 mixRetroflex). In the middle of the continuum, all groups show a learning effect, with more retroflex responses after an exposure that used ambiguous sounds in words with an underlying retroflex.

**Figure 2.**
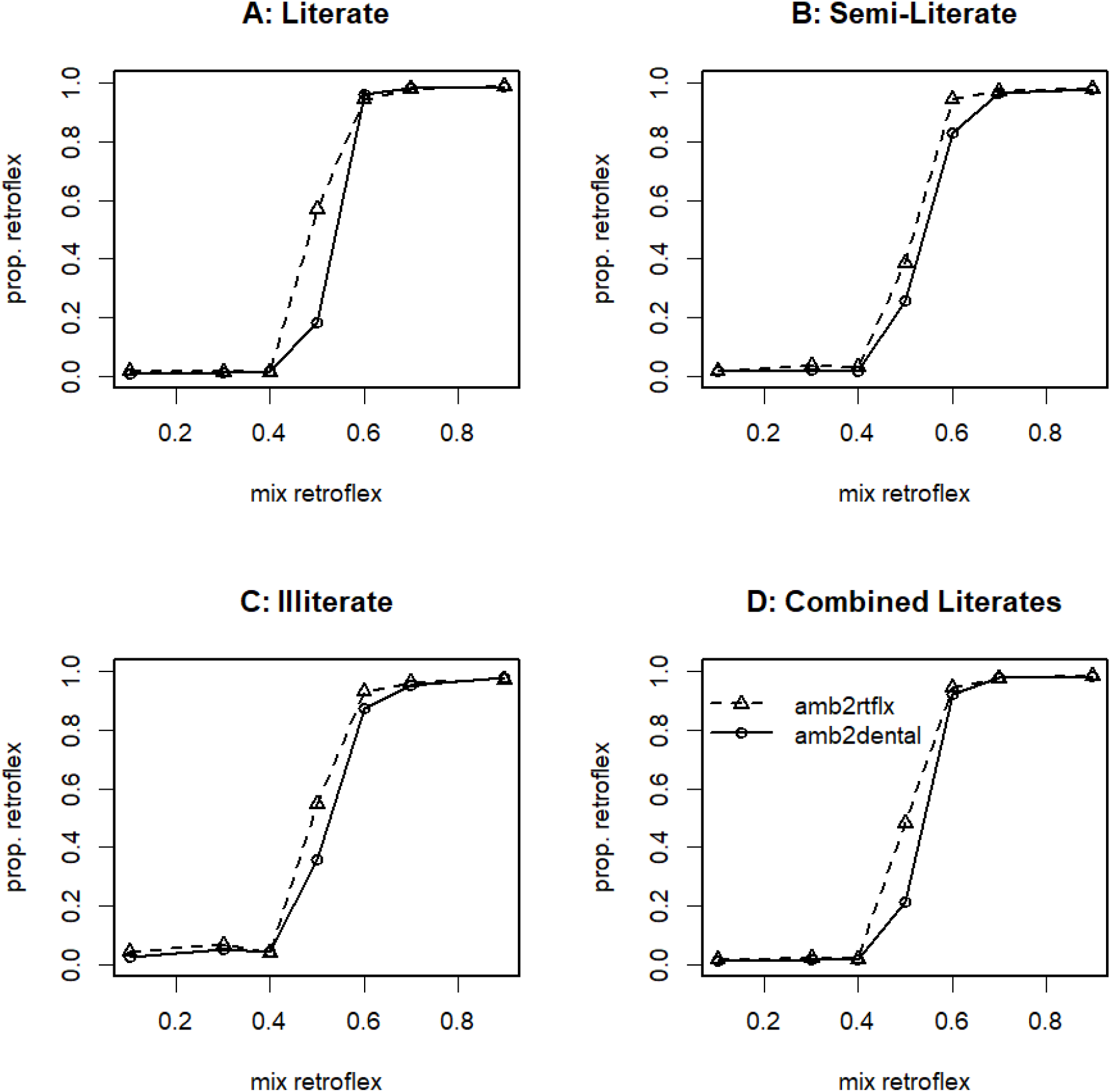
Proportion of retroflex responses during the 2AFC task in the test phase for the literate group (Panel A), the semi-literate group (Panel B) and the illiterate group (Panel C). Panel D shows the data from both groups with reading experience, reflecting the first contrast of the group variable as used in the analysis, literates versus illiterates.

As for the data from the exposure phase, the theoretically critical data from the test phase were analysed with a Bayesian generalized multilevel model with a logit linking function by specifying a Bernoulli family. The statistical analysis focussed on the more ambiguous steps of the continuum, leaving out the guiding stimuli with 10% and 90% proportion of the retroflex signal. For the effect of continuum, the prior was a normal distribution with a standard deviation of ten logit units, as this regression weight can become very large when the identification is categorical. Interactions with this effect of continuum were specified with half of that standard deviation (i.e., Normal(0,5)). All other effects were provided with a prior of Normal(0,2), as above.

The fixed effects were the amount of retroflex stimulus in the mixed signal (or continuum step), the exposure condition, and the reading group. The fixed effects were contrast coded. For Exposure, the group that heard the ambiguous sounds in retroflex words was mapped to 0.5, the group that heard the ambiguous sounds in dental words was mapped to -0.5. The continuum was similarly mapped to the range [-0.5, 0.5], with the most retroflex stimulus mapped to 0.5. For both these effects, there is a strong expectation for a positive regression weight. For the fixed effect of Group, two orthogonal contrasts were used; one contrasting the readers with the illiterates, and one contrasting full literates with semi-literates. Interactions were specified between Exposure and Group, and mixRetroflex and Group. The former might reveal an effect of reading experience on perceptual recalibration, the latter differences in the categoricalness of the identification functions based on reading experience. Participant was a fixed effect, with a random slope for mixRetroflex. Note that all other fixed effects are between participants, hence rendering a random slope over participants inappropriate.

Table 2 shows the results of the analyses. The data indicate that we replicated the overall perceptual learning effect, that is, an effect of Exposure Condition: Participants who heard the ambiguous sound in retroflex contexts during exposure label the ambiguous sounds in the test phase as retroflex more often that participants who heard the ambiguous sound in dental contexts. There also is a (trivial for the purpose of the study) effect of the stimulus continuum on the proportion of retroflex responses; the more retroflex-like the sound, the more retroflex responses. There is strong evidence that the effect of stimulus continuum is moderated by reading experience (i.e., the credible intervals of the interaction exclude zero and the BF is larger than 10) but this is not the case for the interaction of exposure condition by reading experience. For this interaction, the credible intervals clearly include zero and the resulting BF was 0.06, providing strong evidence in favour of the model without the interaction. This supports the hypothesis that perceptual recalibration (that is the effect of exposure) is not moderated by reading experience.

**Table 2.** Results from the Bayesian mixed-effect models on the proportion of retroflex responses in the 2AFC task in the test phase with the regression weight, the credible interval and a Bayes factor indicating evidence for the presence of a term.

|  | B (SE) | 95% CI | Bayes Factor |
| --- | --- | --- | --- |
| Intercept | -0.36 (0.11) | -0.50-0.15 | na |
| Exposure | 0.57 (0.21) | 0.16-0.97 | 4.20 |
| mixRetroflex | 16.7 (1.06) | 14.72-18.88 | >1000 |
| Group |  |  |  |
| Literate vs. Illiterate | -0.36 (0.21) | -0.79-0.06 | 0.08 |
| Full Literate vs. Semi-literate | 0.21 (0.26) | -0.32-0.73 |  |
| Group : Exposure |  |  |  |
| [Literate vs. Illiterate] : Exposure | 0.24 (0.41) | -0.57-1.04 | 0.06 |
| [Full Literate vs. Semi-literate] : Exposure | 0.04 (0.51) | -0.97-1.03 |  |
| Group : mixRetroflex |  |  |  |
| [Literate vs. Illiterate] : mixRetroflex | 5.87 (1.82) | 2.27-9.42 | 443.73 |
| [Full Literate vs. Semi-literate] : mixRetroflex | 6.00 (2.25) | 1.56-10.39 |  |

Note: Bayes Factors are based on model comparison of a full model with a model minus the effect in question. For interactions, the comparison model is the full model with all effects and for main effects, a model with all main effects was compared to one with the effect in question removed.

## Discussion

Spoken-word recognition is typically assumed to involve processing small ‘sound’ segments such as phonemes (e.g., McClelland & Elman, 1986; Norris, 1994). If such segment-sized units are functional in spoken-word recognition, then learning to read (i.e. learning to map ‘orthographic segments onto speech’) may change spoken-word recognition by changing the phonological representations of words (Kolinsky et al., 2021). In other words it is conceivable that orthographic experience directly shapes the representational structure of the spoken word-recognition system. Past behavioral research using low-level speech perception tasks did not find clear performance differences between literate and illiterate adults (despite that both groups typically differ substantially in a great number of other cognitive tasks) but it has recently been suggested that reading experience leads to *more precise* pre-lexical phonological representations (Kolinsky et al., 2021).

We set out to test whether reading experience changes the fidelity of pre-lexical phonological representations in the Tamil language, which is written using an abugida writing system that explicitly codes small (‘phoneme-like’) speech segments. To that end, we performed a perceptual learning experiment with three participant groups that differed in their reading ability (literates, semi-literates, and complete illiterates). The perceptual learning paradigm has previously been found to be particularly useful to investigate prelexical phonological representations (McQueen et al., 2006; Mitterer et al., 2013, 2018).

We observed a clear perceptual learning effect in all groups. Importantly, for the purpose of our study we found that completely illiterate participants performed as well as literate participants in the perceptual learning task. The performance of the illiterates in the perceptual learning task is particularly exceptional given that illiterate participants perform less well on almost any cognitive task that is administered to them (Morais & Kolinsky, 2019; Huettig & Mishra, 2014), including the working memory and Raven’s tasks administered in the present study. In short, despite low scores in working memory and non-verbal intelligence we observed remarkable perceptual learning capabilities in our illiterate participants. The use of the perceptual learning paradigm, the careful design, and large numbers of participants with varying literacy levels leads us to conclude with considerable confidence that pre-lexical phonological representations are not shaped by Tamil reading experience. This interpretation of the results receives additional strong support from the Bayesian analyses we carried out. Learning to read Tamil thus does not appear to restructure phonological representations.

### Alternative interpretations

It is noteworthy though that reading experience led to a difference in the steepness of the categorization curves in our study. What explains this difference? Our study replicates in this regard the findings of Kolinsky et al. (2021) who found a similar reading-related difference in the steepness of categorization. The two findings however point in opposite directions, with the perceptual-learning data suggesting that reading experience does not influence pre-lexical phonological representations while the categorical-perception data suggest the opposite. We conjecture that the perceptual-learning data are more compelling since they show a completely implicit use of pre-lexical representations from exposure to test and is found with explicit measures (such as the 2AFC task) as well as with implicit measures (such as cross-modal priming, McQueen et al., 2006; Sjerps & McQueen, 2010). It is hence very likely that the explicit component of the 2AFC task is the source of the steepness difference between literate and illiterate participants. Note that it is a very common observation in 2AFC tasks that participants sometimes mention that they “pressed the wrong button”. Participants say that they heard, for instance, “ban” but pressed the button that would be associated with a “pan” response. Given that there is a clear verbal-working memory component in such tasks, and verbal working memory is strongly influenced by literacy (Demoulin & Kolinsky, 2016; Smalle et al., 2019), it is likely that differences in the steepness of identification curves are a result of such errors in task performance. This account is supported by the finding that the categorization functions of illiterates often fail to reach 0% and 100%, that is, they still press the “wrong” button even if the acoustic input is unambiguous. These failures of categorization function to have a range from 0% to 100% are obvious in the performance data of illiterates in this study as well as in earlier studies (Kolinsky et al., 2021; Serniclaes et al., 2005).

In the present study, all our participants were speakers of Tamil. Most previous studies investigating effects of learning to read on phonological restructuring were carried out with alphabetic scripts (e.g., French, Portuguese). Tamil script in contrast is an abugida writing system of the Indic type (Daniels, 2021). Are effects of learning to read on phonological restructuring language- and/or script-specific? This is a logical possibility, which receives some support from a recent neuroimaging study carried out with Hindi speakers of varying literacy levels (Hervais-Adelman et al, 2021). On the other hand, there are good reasons that speak against the interpretation that the absence of an effect of learning to read on pre-lexical phonological representations is purely language- and/or script-specific.

As described in the introduction, sounds in Tamil script are represented at *both* the syllable and phoneme level. All consonants can be written as single elements and most syllables are combinations of separate vowel and consonant elements. As such, the Tamil script offers much more evidence for segments than, for instance, the relatively opaque English orthography does. This makes us quite skeptical about suggestions that our results do not extent to alphabetic scripts such as English but further research could usefully be conducted to address this question.

### Final theoretical considerations

Our present findings suggest that the field needs to take some theoretical challenges (for the notion that learning to read results in phonological restructuring) more serious. First, it is important to realize that phonological restructuring would result in similar grain sizes of pre-lexical representations in the visual and auditory domain. Morais and Kolinsky (2019) have correctly pointed out in this regard how problematic the assumption is (often made by reading instructors) that letters stand for speech sounds. Indeed, there is a consensus among researchers that any relation is far more complex. The suggestion that graphemes (rather than letters) relate to phonemes, however, is a very problematic because there is strong evidence that pre-lexical phonological representations are not phonemic (Dahan & Mead, 2010; Mitterer et al., 2013, 2016, 2018), but instead stand for “speech sounds” or (allo-)phones. Establishing *pre-lexical correspondences* between graphemes and the more speech-sound based representations used in spoken-word recognition are thus a much more difficult problem than is typically acknowledged. In the absence of a straightforward one-to-one grapheme-phoneme correspondence, the same grapheme would have to map onto many phonological units (e.g., the grapheme ‘t’ in English needs to map onto an aspirated stop, an unaspirated stop, an unreleased word-final stop, and a flap), and sometimes, the same speech sound may map on different graphemes even in an otherwise shallow orthography, such as German (e.g., [x] maps on the graphemes ‘ch’ and ‘g’ in word-final position, see Mitterer & Müsseler, 2013).

Second, there is the distinction that is typically made between (potential) online recruitment of orthographic representations during speech perception and (potential) reading-related phonological restructuring (e.g., Dehaene et al., 2010). According to the online recruitment account, orthographic information is recruited early in spoken-word recognition. Even before lexical access, incoming phonemes activate (corresponding) graphemes (Frost & Ziegler, 2007; Pattamadilok et al., 2014; Perre et al., 2009; Salverda & Tanenhaus, 2010). This is seen as an inevitable consequence of an interactive processing system, which learns to associate graphemes with phonemes during the process of learning to read (Ziegler & Ferrand, 1998). A consequence of this feedback loop is that lexical access is faster for words in which the phoneme-grapheme relations are consistent, but slower for words with inconsistent relations. We acknowledge explicitly here that our study addressed ‘direct’ phonological restructuring only. We draw attention here however to the problem that the same issue that makes phonological restructuring unlikely (the mismatch in grain size between pre-lexical phonological representations and graphemic representations), also poses a problem for the idea of *routine* online recruitment of orthographic representations during spoken-word recognition: links between speech-based and orthographical representations are difficult to establish if they have different (and depending on context very inconsistent) grain sizes.

Third, while phonological restructuring and orthographic recruitment in spoken-word recognition are independent hypothesis (i.e., both could be correct or false), they are intertwined in ways that has not figured prominently in discussions in the literature. For pre-lexical phonological representations to activate orthographic representations, they must be commensurable. That is, if spoken-word recognition would be based on a syllabic code, for instance, while visual-word recognition would rely on graphemes, it would be difficult to see how they could activate each other. Phonological restructuring would therefore facilitate orthographic recruitment by equalizing the grain size of pre-lexical representations.

Finally, given the complexity of this mapping problem, we conjecture that our findings have clear implications for the debate about the degree of interactivity between different levels of representations during human information processing. More often than not, this question has been cast in terms of competing paradigms, contrasting an interactive view with a modular view of the mind (Ziegler, 2018). We suggest that it might be more beneficial to focus on a functional perspective instead. Rather than assuming modularity or interaction for modularity/interaction’s sake, we believe that it would be useful for the field to consider more thoroughly whether and when interaction is actually beneficial for the language user in a given situation. A functional theoretical perspective can straightforwardly explain robust effects of visual-on spoken-word processing, as found for instance in predictive language processing (Huettig & Pickering, 2019; Mani & Huettig, 2014; Mishra et al., 2012) or transfer of word learning between the modalities (Bakker et al., 2014). In these cases, it is helpful for knowledge and processes to be transferred between modalities. For the purposes of word recognition such transfer however is counterproductive.

## Footnotes

1 Jesse and McQueen (2011) found that word-initial fricatives cannot trigger learning in a lexical decision task, potentially because participants can only infer the intended fricative some time after hearing it. With a picture verification task, this is not the case. Seeing a picture of a flamingo can be sufficient for participants to infer the intended fricative when hearing [^s^/_f_la..].

2 It is important to realize that the range of the predictors is important for understanding the implications of the priors. If the range of the predictors is from -1 to 1, the regressions weights will have half the values compared to predictors ranging from -0.5 to 0.5.

